# Salt-induced first-order structural transition in a DNA-interacting protein

**DOI:** 10.1101/2021.04.15.439963

**Authors:** Yue Lu, Ying Lu, Jianbing Ma, Jinghua Li, Xingyuan Huang, Qi Jia, Dongfei Ma, Ming Liu, Hao Zhang, Xuan Yu, Shuxin Hu, Yunliang Li, Chunhua Xu, Ming Li

**Affiliations:** Beijing National Laboratory for Condensed Matter Physics, Institute of Physics, Chinese Academy of Sciences, Beijing 100190, China; Songshan Lake Materials Laboratory, Dongguan, Guangdong 523808, China; University of Chinese Academy of Sciences, Beijing 100049, China

**Author notes:** These authors contributed equally to this work (Yue Lu, Ying Lu, Jianbing Ma).

## Abstract

Thermodynamics and structural transitions on protein surfaces remain relatively understudied and poorly understood. Wrapping of DNA on proteins provides a paradigm for studying protein surfaces. We used magnetic tweezers to investigate a prototypical DNA-interacting protein, i.e., the single-stranded DNA binding protein (SSB). SSB binds DNA with distinct binding modes the mechanism of which is still elusive. The measured thermodynamic parameters relevant to the SSB-DNA complex are salt-dependent and discontinuous at the bind-mode transitions. Our data indicate that free SSB undergoes salt-induced first-order structural transitions. The conclusion was supported by the infrared spectroscopy of SSB in salt solutions. Ultrafast infrared spectroscopy further suggests that the transitions are correlated with surface salt bridges. Our work not only unravels a long-standing mystery of the different binding site sizes of SSB, but also would inspire interests in thermodynamics of protein surfaces.

---

Wrapping of DNA oligomers on protein surfaces is critical to many DNA transactions [1]. In general, a protein that wraps DNA has a large number of cationic and anionic sidechains in the vicinity of wrapping interfaces. Many cationic groups on the protein surface that form ion pairs with DNA phosphates can form dehydrated surface salt bridges (hydrogen-bonded ion pairs) with neighboring anionic sidechains [1,2]. Many salt bridges are required to create a cationic surface complementary to the anionic DNA phosphates. Patterns defined by the salt bridges therefore dictate the path of DNA wrapping. For instance, mutation of Asp26 to Ala in the histone-like HU proteins in *Bacillus subtilis* (HBsu) increases the DNA site size for HBsu from 10-13 base pairs to >25 base pairs because the surface exposure of Lys3 induced by the substitution of its salt-bridging partner resulted in the enhanced DNA compaction [3]. Despite intense investigations, thermodynamics of salt bridges on protein surfaces has remained incomplete. Binding of SSB to single-stranded DNA (ssDNA) provides a paradigm for studying protein surfaces [4–6]. *E. coli* SSB forms stable homo-tetramers at sub-nanomolar concentrations [7,8] whose four DNA binding domains enable SSB to bind ssDNA in a variety of binding site sizes [4,9,10] [**Figure 1(a)**]. Thermodynamic studies indicated that electrostatic interactions play a major role in SSB–ssDNA binding [11,12]. Fluorescent analysis of the binding site size as a function of salt concentration displayed sigmoidal features with distinct plateau regions that differ in site size, namely 35, 56 and 65 nucleotides (nt) per bound SSB tetramer, referred to as (SSB)_35_, (SSB)_56_ and (SSB)_65_, respectively [10]. Although the transitions attracted much attention since the phenomenon was discovered a few decades ago [13–16], the underlying mechanism is still elusive. Herein, we use magnetic tweezers (MT) and infrared spectroscopy to investigate the thermodynamics of SSB binding to ssDNA.

**FIG. 1.**
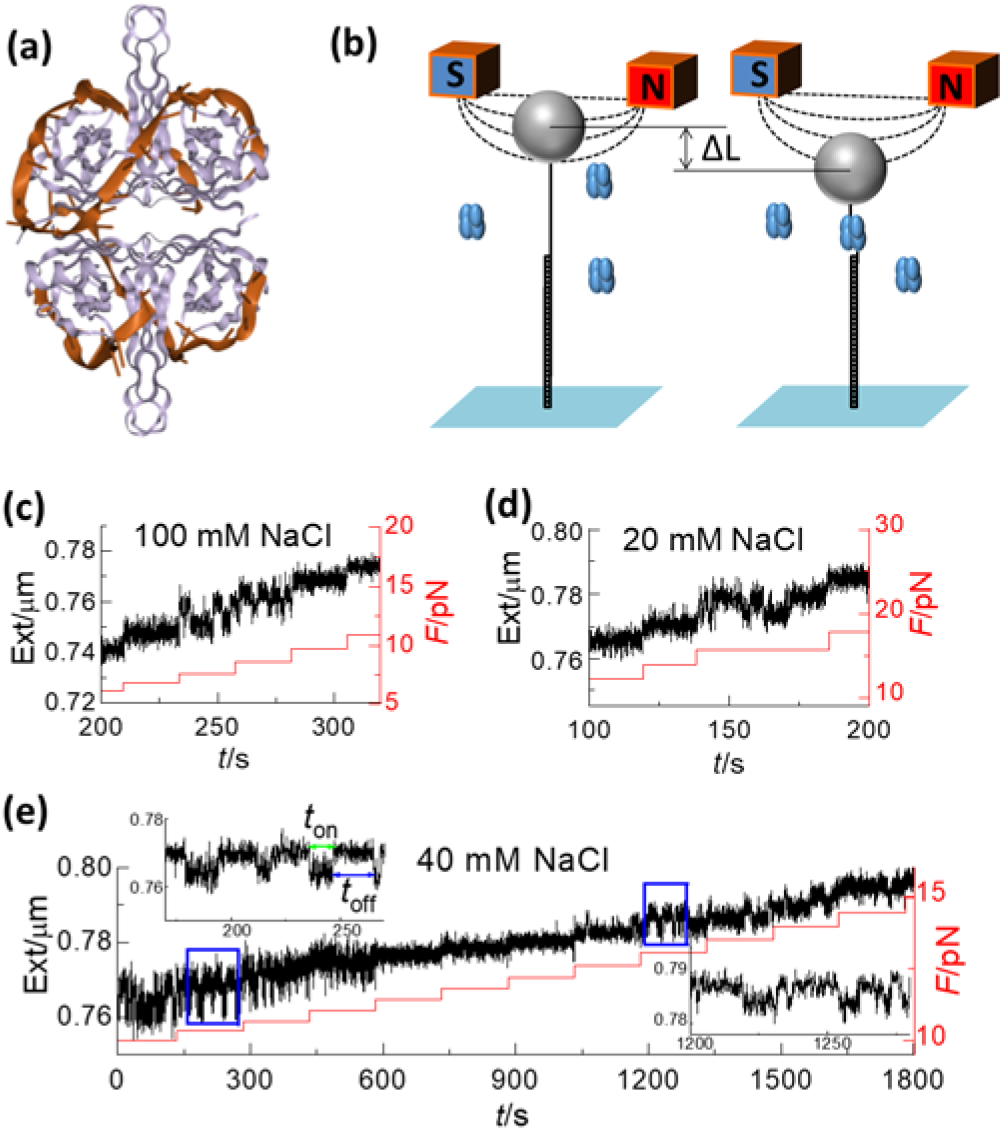
Magnetic tweezers assay of the binding/unbinding kinetics. (a) Structure of (SSB)_65_ (Protein Data Bank ID number 1EYG). (b) Schematic diagram of the experiments. (c-e) Typical time traces of the DNA extension (black lines) at different forces (red lines). Insets in (e) represent details in the blue rectangles, [SSB] = 10 nM.

## MT analysis of SSB binding to 70 nt ssDNA

Binding free energy is the most frequently used parameter to quantify bimolecular interactions and is often calculated by

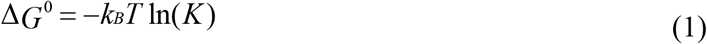

where *k*_B_ is the Boltzmann constant, *T* the temperature and *K* the intrinsic equilibrium reaction constant for a ligand *L* binding to a substrate *S* to form a complex *LS*,

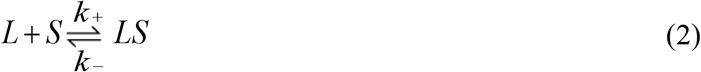

In Eq. (2), *k*^+^ and *k*^−^ are the binding and unbinding rate, respectively, from which one can calculate the reaction constant *K*=*k*^+^/*k*^−^. It is often challenging to measure *k*_0_ accurately in ensemble-averaging experiments if *K* is extremely large. It becomes even more challenging when competitive binding modes with different rates must be discerned. We used single-molecule techniques to resolve the challenge. The different binding modes can be readily distinguished when the measurements of *K* are performed at single-molecule level [17–19]. In addition, the reaction kinetics can be regulated by force because a reaction usually results in configurational changes [20–22]. When force is added slowly enough so that the system can remain quasi-static, the (observed) bimolecular reaction constant changes to

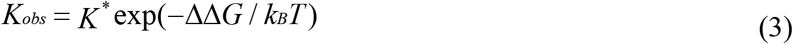

where

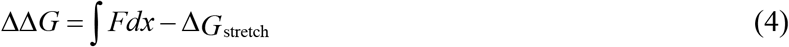

and *K*^*^=[*L*]*k*^+^/*k*^−^ is the pseudo-rate constant [23]. The pseudo-rate constant is proportional to ligand concentration [*L*] because, in single-molecule assays, one usually considers the probability of finding a substrate *S* in a certain state rather than its concentration [23]. The integration in Eq. (4) represents the work done by the force and Δ*G*_stretch_ takes into account the fact that the force not only changes the equilibrium, but also increases the elastic energy of the system [20,22]. When a critical force is exerted so that *K*_obs_ = 1, the binding free energy Δ*G*^0^ is equal to −ΔΔ*G* and can be readily calculated using Eq. (4).

As illustrated in **Figure 1(b)**, we connected one end of a 70-nt ssDNA to a magnetic bead and the other end to a dsDNA which is in turn tethered to the bottom of the fluidic chamber. The binding of SSB to ssDNA resulted in reduction of the end-to-end distance of the DNA. When the force was increased to a certain value, the ssDNA became unraveled, resulting in increase of the end-to-end distance. Binding of a new SSB reduces the distance again. The distance changed repeatedly when the binding and unbinding occurred repeatedly [**Figures 1(c)** to **1(e)**]. The critical force, *F*_c_, at which the binding/unbinding is balanced, depends on the salt concentration. For example, *F*_c_ ≈ 8 pN when [NaCl] = 100 mM [**Figure 1(c)**], whereas *F*_c_ ≈ 16 pN when [NaCl] = 20 mM [**Figure 1(d)**]. Interestingly, at an intermediate salt concentration, e.g., 40 mM, we observed two critical forces, one at ~10 pN and the other at ~14 pN [**Figure 1(e)**].

The kinetics of binding and unbinding can be derived from the time traces of the end-to-end distance, with the large value representing the unbound state and the small value the bound state [Insets in **Figure 1(e)** and **Supplementary Fig. 1**] [24]. We recorded dwell times (*t*_on_ and *t*_off_) of both the bound and unbound states, the histograms of which display exponential decays [**Supplementary Fig. 1**] [24]. Inverses of the characteristic times are the (observed) binding and unbinding rates, respectively. The force dependences of the binding/unbinding rates thus obtained are displayed in Supplementary Fig. 2(a) [24]. The binding rate decreases with force, whereas the unbind rate increases with force. The critical force *F*_c_ can therefore be readily determined from the crossover of the two force-dependent curves. To exclude the possibility that the reduction in distance is due to the rewrapping of the same single SSB, we varied [SSB] and found that the binding rate increased linearly with [SSB], while the unbinding rate was independent of it [**Supplementary Fig. 2(b)**] [24].

## Discontinuousness of thermodynamic parameters

We set out to assess thermodynamic parameters relevant to the bimolecular reaction. They are the critical force *F*_c_, the rate at balance *kF*_c_ and the wrapping length Δ*X* of ssDNA on SSB [**Figure 2**]. Our data showed that SSB binds to ssDNA with two different modes in the two regions studied. Overall, the critical force decreases with the increasing of the salt concentration, indicating that the DNA-protein interaction strength reduces when the salt concentration increases [**Figure 2(a)**] [11,29]. However, *F*_c_ jumps down near the transition concentration. In accordance with this, *kF*_c_ and Δ*X* are also discontinuous. We converted the wrapping length [**Figure 2(c)**] to the binding site sizes [**Figure 2(d)**] according to the structures of the SSB-ssDNA complexes [30,31]. The resulting site sizes are about 17 and 35 nt, respectively in the two [NaCl] ranges studied, which were much shorter than expected. We used FRET to check the site sizes of the SSB-ssDNA complex in various [NaCl] [**Supplementary Fig. 3**] [24]. The resulting FRET values are 0.65 and 0.4 in 10 and 100 mM NaCl solutions, which are in accordance with the site sizes of 35 and 56 nt, respectively. Our MT results therefor suggested that, depending on the binding mode, a different number of nucleotides are more susceptible to tension-induced unraveling, indicating that the ssDNA segments associated with the four monomers are not necessarily equivalent [31,32]. This is consistent with the negative cooperativity observed by Lohman et al., who showed that the wrapping free energy landscape of (SSB)_65_ is not distributed evenly along the nucleotide [31].

**FIG. 2.**
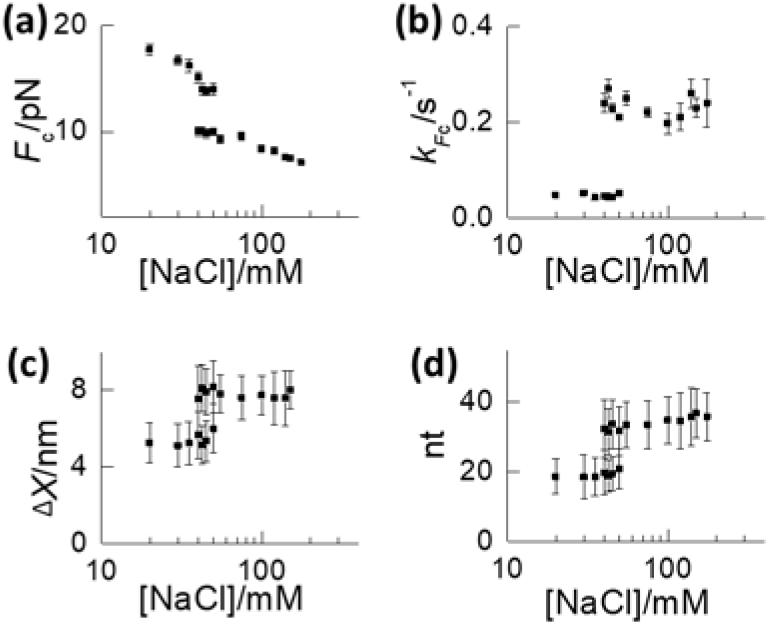
Thermodynamic parameters versus salt concentration for SSB binding to 70 nt ssDNA. (a) Critical forces. (b) Binding rates at critical forces. (c) Binding lengths. (d) Binding site sizes. The error bars in (a) are got by force measured fluctuation; errors bar in (b) ~ (d) are got by fitting error.

We noticed that the force dependence of the binding rate is different from that of the unbinding rate near the critical force [**Supplementary Fig. 2(a)**] [24]. This enabled us to calculate the positions of the free energy barrier of both the forward and the backward reactions through

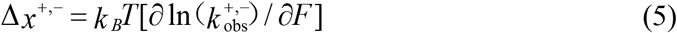

where Δ*x^+^ = x*_barrier_ - *x*_start_ and *Δx*^−^ = *x*_end_ - *x*_barrier_ [**Supplementary Fig. 4**] [24]. The positions of the barriers in the free energy landscapes [**Supplementary Fig. 5**] [24] at various salt concentrations are all equivalent to a structure with ~9 nt bound to an SSB tetramer. The result implicates that the initial binding process is critical for the thermodynamics of SSB binding to ssDNA.

## Binding of SSB to 20 nt ssDNA

In general, discontinuousness of thermodynamic parameters may indicate a first-order phase transition. As can be clearly seen from **Figure 2**, in a narrow range of salt concentrations, we obtained two values for each of the parameters. This is in consistence with the results illustrated in **Figure 1(e)** in which we observed two kinds of binding/unbinding events at a single salt concentration with [NaCl]=40 mM (see the two blue rectangles in **Figure 1(e)**). The results imply that two states coexisted near the phase transition point. We expected to obtain the free energy of the reaction to see what underlies the molecular mechanism of the binding mode transitions. In principle, one can use Eq. (4) to calculate the null-force binding free energy Δ*G*^0^, which is equal to −ΔΔ*G* in Eq. (4). However, we were not able to measure accurately the response of the SSB-ssDNA complex at very low forces where a fraction of the ssDNA, ~18 nt in (SSB)_35_ and ~21 nt in (SSB)_56_, was peeled off the protein. It is known that the shortest length of ssDNA that is relevant to the initial binding is about 17 nt [31]. Our results also implicated that the initial binding of the first 17 nt is critical for the thermodynamics of SSB binding to ssDNA [**Supplementary Fig. 4** and **5**] [24]. We therefore performed experiments of SSB binding to 20 nt ssDNA. The binding/unbinding balances are like that for a 70 nt ssDNA [**Figure 3** and **Supplementary Fig. 6**] [24]. Again, the reaction constant *kF*_c_ is discontinuous [**Figure 3(b)**] although the wrapping length is the same for the two binding modes [**Figure 3(c)**]. We used Eq. (4) to calculate the null-force reaction constant, in which the work done by the force is *F*_c_×Δ*X* and the stretching free energy Δ*G*_stretch_ is the one consumed to stretch the 20 nt ssDNA from *F*=0 to *F*=*F*_c_. Surprisingly, the resulting binding free energy as a function of salt concentration is not continuous at the transition point [**Figure 3(d)**]. Neither would the two parts of the free energy curve be extrapolated to intersect in the regions nearby.

**FIG. 3.**
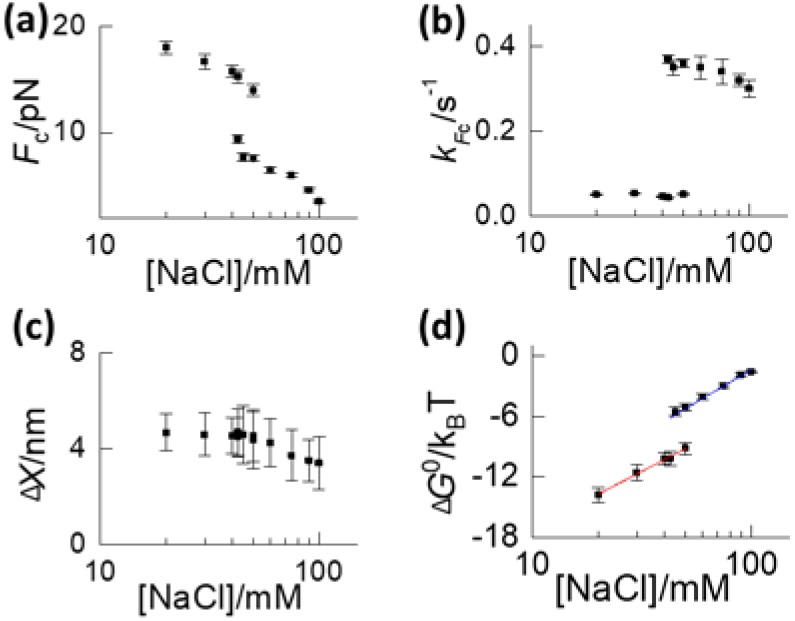
Thermodynamic parameters for SSB binding to 20 nt ssDNA. (a) Critical forces. (b) Binding rates at critical forces. (c) Binding lengths. (d) Binding free energy. The error bars in (a) are got by force measured fluctuation; errors bar in (b) ~ (c) are got by fitting error; error bars in (d) are calculated by Propagation of uncertainty relationship from error of measured force and length.

## Infrared spectra changes of free SSB versus salt concentration

According to thermodynamics, the free energy function of a system undergoing first-order phase transition should be continuous, but its first order derivation is not. This was not the case for the SSB-DNA complex, we therefore wondered if the phase transition is not of the complex, but rather the free SSB itself. To check this idea, we performed Fourier transform infrared (FTIR) spectroscopic analysis of free SSB in different NaCl solutions. At a first glance, the FTIR spectra at different salt concentrations were almost the same [**Supplementary Fig. 7**] [24], suggesting that the globalized conformation of SSB does not change much in different salt solutions. In order to see the spectral differences in details, we decomposed the wide peaks in the spectra by using the second-order derivative and deconvolution method [**Supplementary Fig. 8** and **10**] [24,34]. It turned out that the changes were mainly manifested in the amide I′ window (1600 to 1700 cm^−1^) for backbones and the CH stretch window (2800 to 3000 cm^−1^) for side chains [**Figures 4**]. The wide peak in the amide I′ window can be decomposed nicely into seven Gaussian peaks [**Figure 4(a)**] which can be divided into two groups. While little changes can be noticed for four of them (1624, 1641, 1679 and 1693 cm^−1^), obvious changes can be seen for three peaks at 1650, 1661 and 1670 cm^−1^ [**Figure 4(c)** and **Supplementary Fig. 9**] [24], which corresponds to random coils, α-helices and β-turns [35], respectively. When normalized to their maxima, it is obvious that the components at 1650 and 1670 cm^−1^ jumped down companying with the simultaneous bouncing up of 1661 cm^−1^ when the salt concentration was increased from 12 to 17 mM. This clearly demonstrates that a conformational phase transition occurred with abrupt reduction in random coil and β-turn components and increase in α-helix components. Similarly, the wide peak centered at 2950 cm^−1^ in the CH stretch window can be decomposed into five Gaussian peaks located at 2922, 2931, 2941, 2951 and 2968 cm^−1^, respectively [**Figure 4(b)**]. When normalized to their maxima, the jumps of the peak intensities became more obvious [**Figure 4(d)** and **Supplementary Fig. 11**] [24], which is more consistent with the behaviors of vibrational modes in amide I^’^ window. It is noteworthy that the absolute changes of the peak intensities are very small, implicating that only partial conformations were slightly adjusted while the overall backbone remained almost unchanged.

**FIG. 4.**
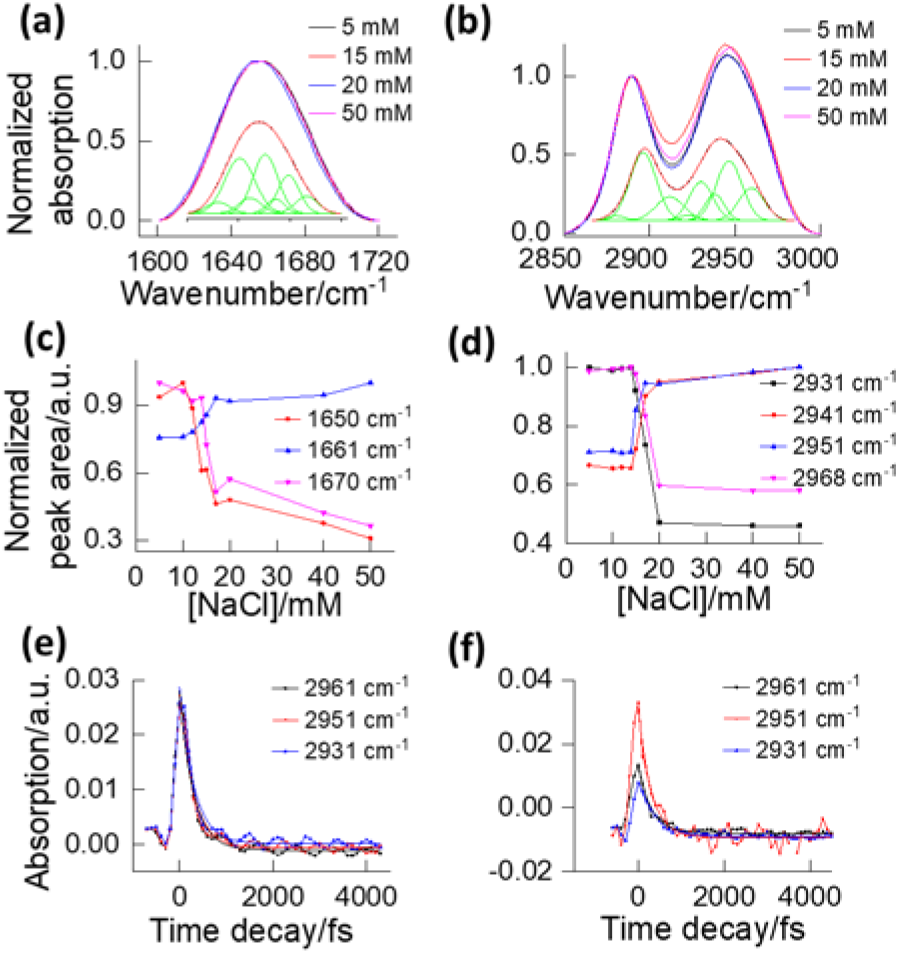
FTIR spectra and vibrational energy relaxation (VER) dynamics of free SSB. (a) FTIR spectra in the amide I′ window (1600 to 1700 cm^−1^). Inset shows the decomposition of a peak. (b) FTIR spectra in the CH stretch window. Inset shows the decomposition of two peaks. (c) Normalized FTIR spectral intensity changes of SSB in the amide I′ window for the vibrational modes at 1650, 1661 and 1670 cm^−1^. (d) Normalized FTIR spectral intensity changes of SSB in the CH stretch window for the vibrational modes at 2931, 2941, 2951 and 2968 cm^−1^. (e) VER dynamics of SSB at 5 mM NaCl. (f) VER dynamics of SSB at 17 mM NaCl.

We measured the vibrational energy relaxation (VER) in the CH stretch window with time-resolved mid-infrared pump-probe spectroscopy to check the dynamics of conformational changes. Three frequencies (2931, 2951 and 2961 cm^−1^) were selected to represent different contributions involved in the averaged signal for side chains [**Figures 4(e)** and **4(f)**]. The time decay constants of the VER at 5 mM NaCl [**Figure 4(e)**] for the three frequencies are 212, 204 and 205 fs, respectively. When the salt concentration was increased to 17 mM [**Figure 4(f)**], the time decay constants became 275, 206 and 207 fs, respectively. Although the decay times did not change much, the maxima of VER at time zero for the three frequencies behaved differently at the two salt concentrations. While they are almost of the same values in the 5 mM NaCl solution [**Figure 4(e)**], the signal for 2931 cm^−1^ and 2961 cm^−1^ are almost two times weaker than that for 2951 cm^−1^ in the 17 mM NaCl solution [**Figure 4(f)**]. These changes would make the spectra narrower and more centered at the vibrational mode at 2951 cm^−1^, implying that the conformation of SSB would be more compacted at higher salt concentrations. The 200 fs time scale of VER is reminiscent of fast hydrogen-bond breaking and reforming, which may occur in the water environment surrounding the CH3 probes and/or in the protein scaffold [36]. The control experiment with BSA, a protein often used as a probe for dynamic responses of water, did not show any spectral changes when the salt concentration changes [**Supplementary Fig. 12**] [24,37]. Therefore, the changes in SSB may not arise from the hydrogen bonds in water but rather from the hydrogen bonds and/or hydrogen-bonded ion pairs (salt bridges) on the protein surface. This is consistent with a recent quantitative analysis of the effects of ionic liquid on protein conformation that also emphasized the impact of the salt bridges [38] which play important role in conformation and stabilities [39]. It was estimated that essentially all 30 lysine, arginine and histidine residues in the vicinity of the SSB-DNA interface can form salt bridges to carboxylates [1,3,40]. We hypothesize that NaCl may induce first-order structural transition of the salt-bridge pattern on the surface of SSB, resulting in different binding modes with polynucleotides.

There exist two populations, (SSB)_35_ and (SSB)_56_, in a wide range of salt concentration (5-50 mM) around the midpoint 17 mM [10]. In our MT assay, the apparent midpoint shifted to ~45 mM, near the upper limit of the transition range. We argue that the shift does not affect our conclusion for the following reasons. The critical force *F*_c_ is higher for (SSB)_35_ (*F*_c_≈16 pN) than for (SSB)_56_ (*F*_c_≈9 pN) [**Figure 2** and **Figure 3**]. At a concentration much lower than 45 mM, when the force is around 9 pN, <u>once a (SSB)_35_ protein already binds the DNA, a (SSB)_56_ protein has no chance to bind the DNA because the force is not large enough to peel off the (SSB)_35_ protein</u>. On the other hand, if a (SSB)_56_ protein binds the DNA first, it is likely to be replaced by a (SSB)_35_ protein very quickly. In both cases, the probability of catching the binding/unbinding of (SSB)_56_ is very low in the MT assay. Only when the salt concentration is increased to near the upper limit of the transition range, the population of (SSB)_56_ is much higher than that of (SSB)_35_ so that the binding/unbinding dynamics of (SSB)_56_ become observable.

In summary, we measure the thermodynamic parameters of SSB binding to ssDNA in a wide range of NaCl concentrations, revealing that an SSB tetramer undergoes structural transition when the salt concentration is increased across the transition point. This may unravel a part of the mystery of the different binding site sizes of SSB. Our discovery left us another mystery: What kind of structural transition it is and what is the mechanism underlying the transition? So far, no evidence has been found in the literature that the main chains of SSB may undergo overall structural transition. Our time-resolved IR measurements implicated that the structural changes might arise from hydrogen bonds and/or salt-bridges on the surface of SSB. Because protein-nucleic acid interactions are common and many of the interactions are electrostatic in nature, we prospect that more proteins will be found to have surface phase transitions. Thermodynamics of protein surfaces deserve certainly further in-depth investigations.

## Acknowledgment

This work was supported by the National Key Research and Development programme of China (Grant No. 2019YFA0709304), National Natural Science Foundation of China (Grant Nos. 11974411 to Ying Lu, and 91753104 to M. Li), Excellent Young Scholars of National Natural Science Foundation of China (Grant No. 12022409 to Ying Lu), CAS Key Research Program of Frontier Sciences (Grant Nos. QYZDJ-SSW-SYS014 to M. Li and ZDBS-LY-SLH015 to Ying Lu), and CAS Youth Innovation Promotion Association (Grant No. 2017015 to Ying Lu). The authors also gratefully acknowledge the support of the K. C. Wong Education Foundation.

