## Supplementary Material for "Salt-induced first-order structural transition in a DNA-interacting protein" (from another version of this preprint: version 1, 2021-04-15)

for

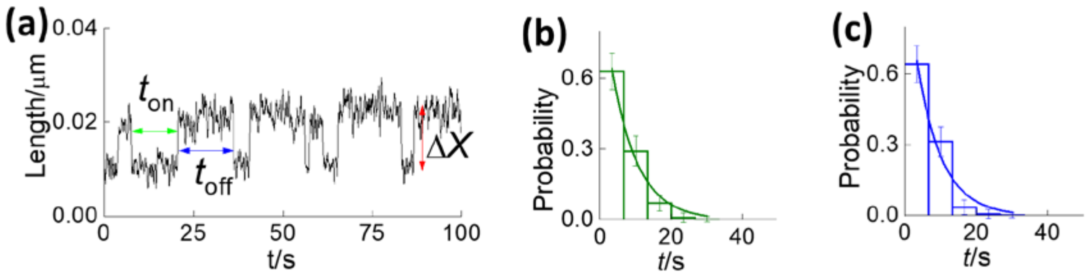


Supplementary Figure 1. The binding/unbinding rates are obtained from the “binding and unbinding” time traces. (a) An example of the binding/unbinding time traces. (b) Dwell time distribution of the binding states. (c) Dwell time distribution of the unbinding states.

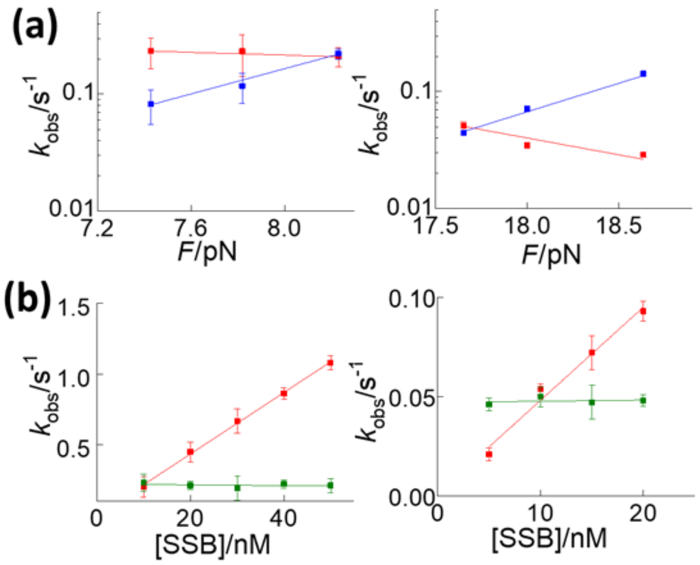


Supplementary Figure 2. The observed binding/unbinding rates. (a) Binding (red)/unbinding (blue) rates vs force at [SSB]=10 nM with [NaCl] = 120 mM in the left panel and [NaCl] = 20 mM in the right. (b) Binding (red)/unbinding (green) rates vs [SSB] at [NaCl] = 120 mM with Force= 8.2 pN in the left panel and [NaCl] = 20 mM with Force= 17.7 pN in the right panel.

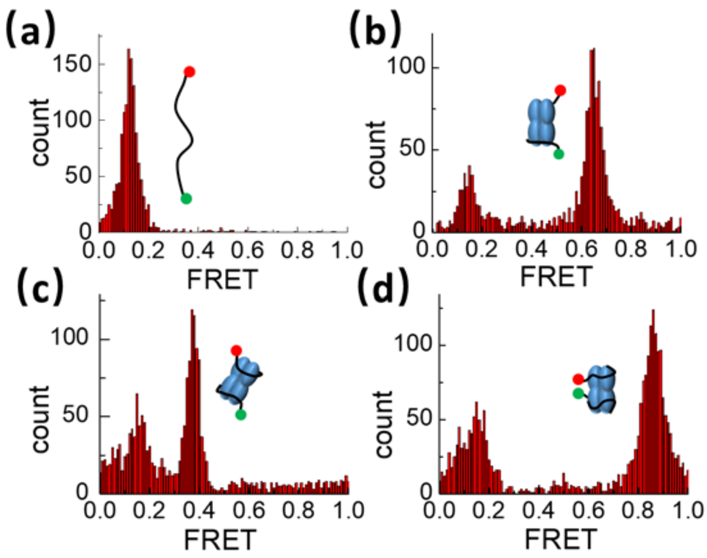


Supplementary Figure 3. Histograms of FRET values of the 70-nt ssDNA and the SSB/ssDNA complexes in different salt solutions. (a) ssDNA alone. (b) SSB/ssDNA in 20 mM NaCl. (c) SSB/ssDNA in 100 mM NaCl. (d) SSB/ssDNA in 300 mM NaCl.

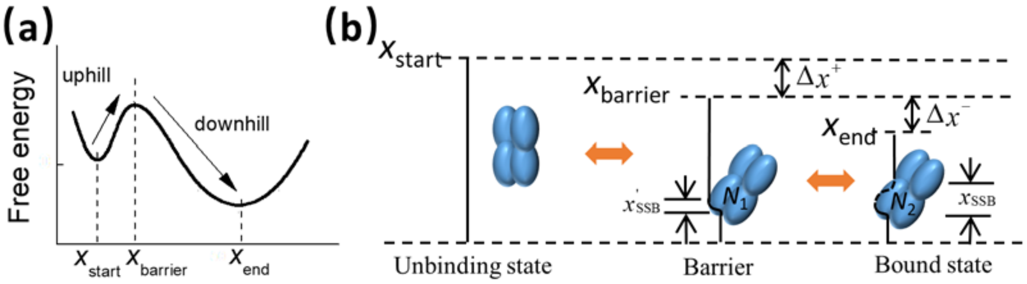


Supplementary Figure 4. Schematic illustration of the SSB/ssDNA bimolecular reaction. (a) The free energy landscape of ssDNA wrapping on SSB. (b) Definition of the lengths used in the calculation by using Eq. (S1), taking $N_2$=17 as an example.

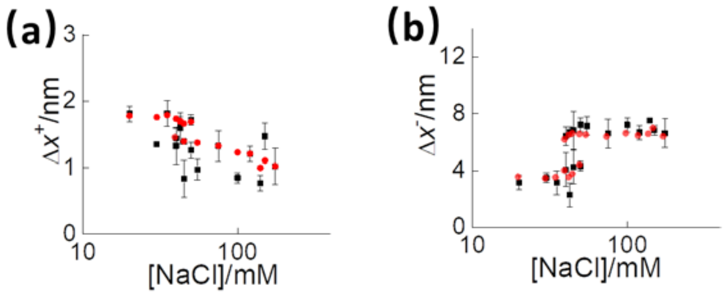


Supplementary Figure 5. Positions of the barrier of the free energy landscape in different [NaCl]. (a) Uphill reaction length $\Delta x^{+}$; (b) Downhill reaction length $\Delta x^{-}$. Black squares are from Figure S2 and Eq. (5). Red dots are from Figure S4(b) and Eq. (S1).

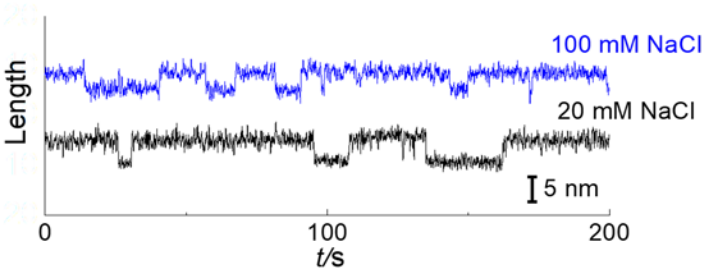


Supplementary Figure 6. Typical binding/unbinding time traces of SSB binding to a 20-nt ssDNA.

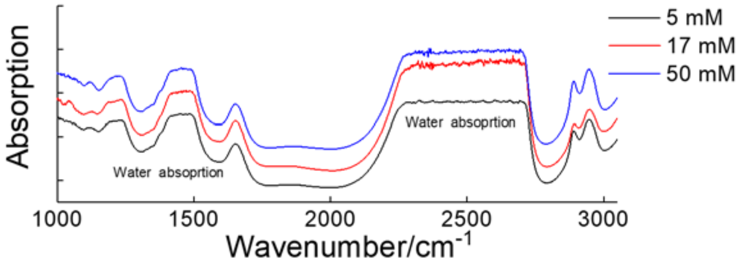


Supplementary Figure 7. Overview of the FTIR spectra of SSB at 5, 17 and 50 mM NaCl in $D_2O$ solutions.

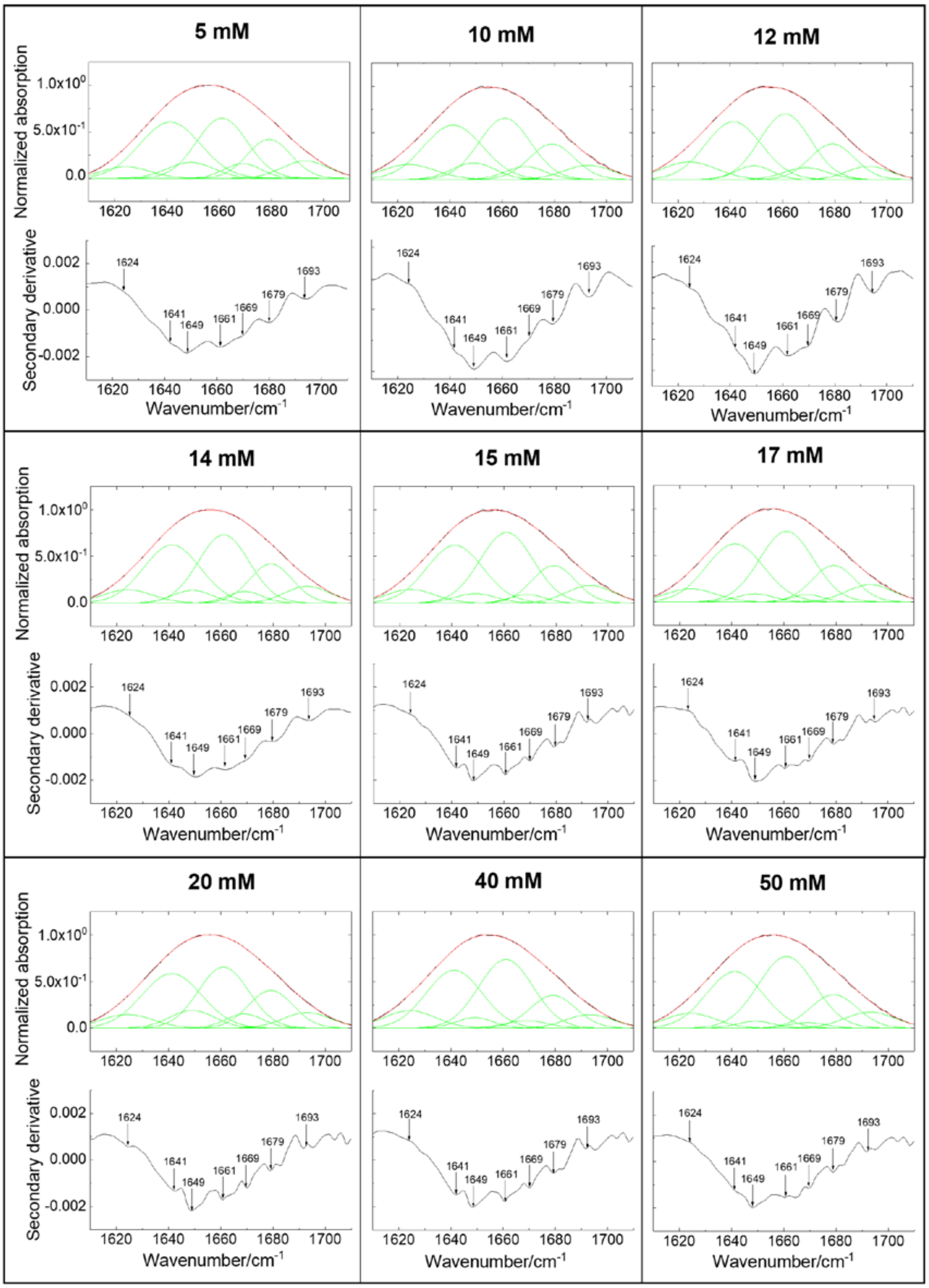


Supplementary Figure 8 The FTIR spectra of SSB in the amide I′ window (1600 to 1700 cm$^{-1}$) at different NaCl concentrations in $D_2O$ and their second derivatives. The decomposed individual absorption bands of the structural components are shown as green lines. The values of the NaCl concentrations are indicated in the figures.

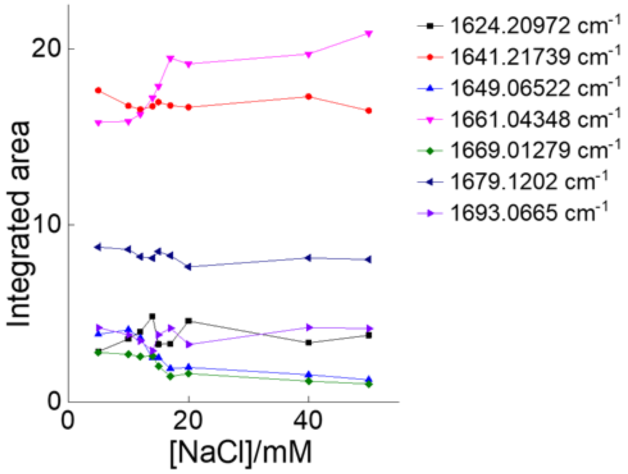


Supplementary Figure 9 Integrated areas of the absorption bands versus salt concentration.

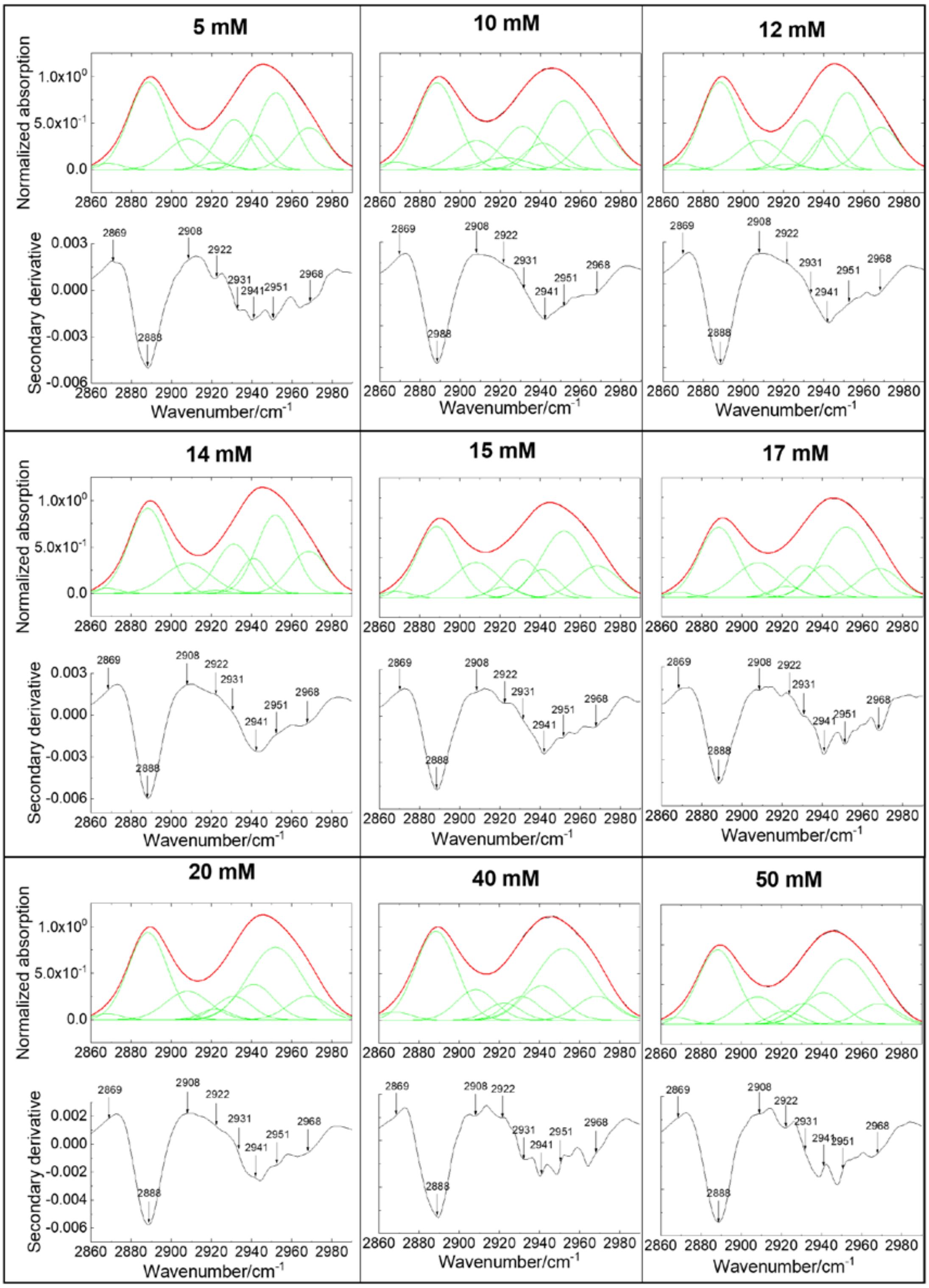


Supplementary Figure 10. The FTIR spectra of SSB in the CH stretch window (2800 to 3000 $cm^{-1}$) in different NaCl concentrations in $D_2O$ and their second derivatives. The decomposed individual absorption bands of the structural components are shown in green lines. The values of the NaCl concentrations are indicated in the figures.

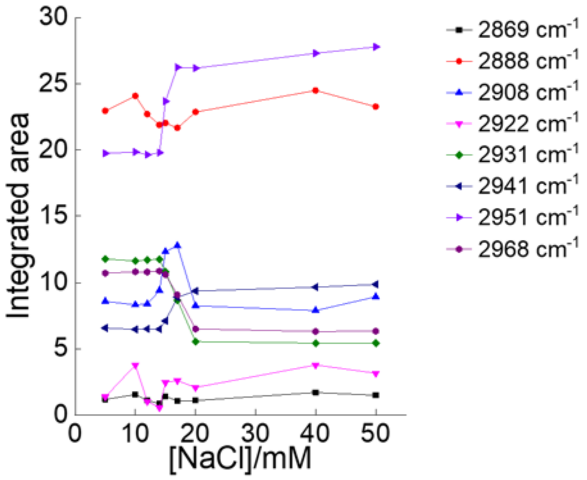


Supplementary Figure 11. Integrated areas of the absorption bands versus salt concentration.

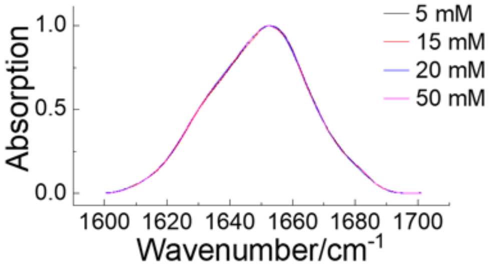


Supplementary Figure 12. FTIR spectra of the BSA protein at 5, 15, 20 and 50 mM NaCl in $D_2O$ solutions.

### *Materials and Methods*

#### DNA constructs and SSB

The DNA constructs for the magnetic tweezers assays contains two parts: a 2300-bp double-stranded DNA as handle and a 70-nt single-stranded DNA (Poly dT) as binding site [1]. The dsDNA handle was synthesized from PCR amplification of the PBR322 plasmid using a 5′-digoxigenin-labeled primer and digested with the Kpn I restriction enzyme (New England Biolabs), leaving a 4-nt 3’ overhang. The ssDNA was purchased from Sangon Biotech (Shanghai, China) with a 5′-biotin modification and annealed with a short oligo to make a double-stranded terminal with a 4-nt 3′ overhang. The double-stranded terminal with a 4-nt 3′ overhang was ligated to the dsDNA handle, resulting in a DNA with one digoxigenin and one biotin on two opposite ends.

The *E. coli.* SSB protein was purchased from Sigma-Aldrich (S3917). SSB is stored in SSB storage buffer (20 mM Tris-HCl, pH 8.0, 0.5 M NaCl, 0.1 mM EDTA, 0.1 mM DTT, 50% glycerol). Before injected into the chamber, SSB was diluted by the reaction buffer (20 mM Tris-HCl, pH 8.0, 20-500 mM NaCl).

#### The magnetic tweezers assay

A flow chamber was made of a coverslip and a slide. The coverslip was thoroughly cleaned and then silanized with Sigmacote (Sigma-Aldrich). Its surface was modified by anti-digoxigenin and blocked by a passivation buffer (10 mg/ml BSA, 1 mM EDTA, 10 mM pH 7.4 phosphate buffer, 10 mg/ml Pluronic F127 surfactant (Sigma-Aldrich), 3 mM $NaN_3$). The flow chamber was placed on an inverted microscope (IX71, Olympus) with an objective (100×, oil immersion, NA 1.45, Olympus). The magnetic beads (2.8 μm diameter, streptavidin coated, Invitrogen) were mixed with DNA substrate in certain proportion; and then injected into the chamber to complete the linkage [2]. Two small NdFeB magnets were mounted above the chamber to pull the DNA-tethered magnetic beads. After checking the single DNA linkage by stretching DNA and observing the length change after twisting, SSB solution was injected and incubated for 10 min. The length of DNA was continuously recorded under different forces.

**FTIR and time-resolved mid-IR spectroscopy**

The FTIR spectra were collected in a Bruker Tensor II FTIR spectrometer by 64 scans at a resolution of 4 $cm^{-1}$.The concentration of SSB was controlled at 4470 μg/ml in $D_2O$ to reduce the water absorption in the amide I′ region, and the proteins in different salt solutions were hold in a sandwich-structure cell with two $CaF_2$ windows separated by a 100 µm path length Teflon spacer. The whole system was sealed and continuously flush out with dry air to reduce the air humidity.

The details of the ultrafast IR pulse generation and signal collection methodologies have been described previously [3]. Briefly, a commercial Ti:sapphire regenerative amplifier (SpectraPhysics, Spitfire) is seeded by a Ti:sapphire oscillator (Spectra Physics, Tsunami) to produce a 1 kHz train of 100 fs pulses centered at 800 nm with an average pulse power of 5 mJ. Approximately 3.5 mJ of the output was used to pump a commercially automated optical parametric amplifier (TOPAS Prime). The generated signal and idler pulses were then used to pump a collinear difference frequency generation module. Finally, the 100 fs mid-IR pulses centered at 2950 $cm^{-1}$ with 20 μJ output energy was obtained. About 5% of the mid-IR radiation was split off to become the probe beam by a 3$^{o}$ $CaF_2$ wedge. The remaining mid-IR became the pump beam and was fed into a mechanical translation stage (IMS600LM, Newport) for time delay. The pump and probe beams were focused onto the sample by a parabolic mirror with the spot size of 80 μm and 120 μm, respectively. The produced signal were collected by the 64-channel detector (FPAS-0144, Infrared Systems Development) combining with 300 mm focal length spectrometer (SP2300i，Princeton instrument).

***Data analysis***

**The binding and unbinding rates**

The kinetics of binding and unbinding can be derived from the time traces of the end-to-end distance, with the large value representing the unbound state and the small value the bound state [Insets in **Figure 1(e)** and **Supplementary Fig. 1(a)**]. The binding and unbinding rates were obtained by analyzing the dwell times ($t_{on}$ and $t_{off}$) of the bound state and unbound state displayed in **Supplementary Fig. 1** (see also **Figure 1**). The histograms of the dwell times of the both states can be fitted to single exponential functions [**Supplementary Fig. 1(b)** and **(c)**]. The inverses of them are the binding and unbinding rates are displayed in **Figure 2** in the main text.

**Force and SSB concentration dependence of the binding and unbinding rates**

The force dependences of the binding/unbinding rates obtained above are displayed in **Supplementary Fig. 2(a)**. The binding rate decreases with force, whereas the unbind rate increases with force. The critical force $F_c$ can therefore be readily determined from the crossover of the two force-dependent curves. To exclude the possibility that the reduction in distance is due to the rewrapping of the same SSB, we varied [SSB] and found that the binding rate increased linearly with [SSB], while the unbinding rate was independent of it [**Supplementary Fig. 2(b)**].

**The full binding site sizes measured in the FRET assay**

We labeled Cy3 and Cy5 onto the two ends of 70-nt ssDNA molecules and tethered it to the surface of the coverslip by the interaction between biotin and streptavidin, and then measured the end-to-end distance by FRET under different salt conditions [**Supplementary Fig. 3**]. The statistical distribution of FRET with a peak at 0.15 displayed in **Supplementary Fig. 3(a)** represents ssDNA without SSB. When SSB was added into the chamber, the peak moved to 0.65 in 20 mM NaCl [**Supplementary Fig. 3(b)**], 0.4 in 100 mM NaCl [**Supplementary Fig. 3(c)**] and 0.9 in 300 mM NaCl [**Supplementary Fig. 3(d)**], respectively. These values can be attributed to the 35-nt, 56-nt and 65-nt binding mode, respectively.

**The barrier of the free energy landscape**

The position of the free energy barrier [**Supplementary Fig. 4(a)**] of both the forward and the backward reactions can be obtained from the data in **Supplementary Fig. 2(a)** by using Eq. (5) in the main text. The resulting $\Delta x^+ = x_{\mathrm{barrier}} - x_{\mathrm{start}}$ and $\Delta x^- = x_{\mathrm{end}} - x_{\mathrm{barrier}}$ are displayed in **Supplementary Fig. 5** (black squares). By definition [**Supplementary Fig. 4(b)**], they are related to the number of bound nucleotides, $N_1$ and $N_2$, and the distance between the two terminals of the bound DNA on SSB, $x'_{\mathrm{SSB}}$ and $x_{\mathrm{SSB}}$

$$\Delta x^+ = N_1 L_{\mathrm{nt}} - x'_{\mathrm{SSB}} \quad \text{(S1a)}$$

$$\Delta x = N_2 L_{\mathrm{nt}} - x_{SSB} \quad \text{(S1b)}$$

$$\Delta x^{-} = \Delta x - \Delta x^{+} \quad \text{(S1c)}$$

where $N_1$ and $x'_{SSB}$ are the number of nucleotides and the corresponding length of the ssDNA segment bound on the SSB before the energy barrier; $N_2$ and $x_{SSB}$ are the number of nucleotides and the corresponding length of the ssDNA upon fully binding the SSB. The length $x_{SSB}$ can be deduced from the structure of SSB-ssDNA complexes. For example, $x_{SSB}$ = 3.5 nm for the $N_2$=17 nt binding mode and $x_{SSB}$ = 5.5 nm for the $N_2$=35 nt binding mode. Taking $N_1 \approx 9$ nt and obtain the corresponding $x'_{SSB} \approx 2.2$ nm from the crystal structure [4,5], we calculated $\Delta x^{+}$ and $\Delta x^{-}$ according to Eq. (S1) and illustrated them in **Supplementary Fig. 5** (red dots). The two sets of data (the black squares from Eq. (5) and the red dots from Eq. (S1)) agree well with each other. Therefore, the barriers of the free energies are at ~9 nt for the both binding modes. Note that, in Eq. (S1), $L_{nt}$ is the size of one nucleotide in the salt solution, which is dependent on the salt concentration as well as the force exerted on the ssDNA. $L_{nt}$ can be calculated according to Saleh *et al* [6].

**Binding of SSB to 20 nt ssDNA**

We also analyzed the binding of SSB to a 20 nt ssDNA. We replaced the 70-nt ssDNA substrate with the 20 nt ssDNA and kept other conditions same in the magnetic tweezers assay. The results were similar to that with the 70 nt ssDNA. The DNA length jumped repeatedly near critical forces in different salt concentrations [**Supplementary Fig. 6**].

**FTIR spectra analysis**

The FTIR spectra of SSB in the range 1000-3050 cm$^{-1}$ were collected at 4 cm$^{-1}$ resolution [**Supplementary Fig. 7**]. The components of the FTIR spectra in the amide I′ window (1600 to 1700 cm$^{-1}$) and the CH stretch window (2800 to 3000 cm$^{-1}$) were further analyzed by using the second-order derivative and deconvolution method [**Supplementary Fig. 8** and **10**]. The curve fitting of original spectrum is performed by stepwise iterative adjustment to determine the shape and position. Firstly, the frequency position of each decovoluted peak under the envelope curve were found with its second derivative. Then the peak positions were fixed, and peak the intensity and width were obtained by fitting the envelope curve. Finally all the variables were iterated to converge with minimum root-mean-square error toward 10$^{-6}$. The assignments of the absorption peaks to the secondary structure elements were based on other studies in which infrared spectra were analyzed for proteins of known secondary structure [7]. The integrated area of the individual peak was extracted to represent the absorption band for each secondary structure element. The component of each secondary structure element versus salt concentration were plotted to check the changes of protein conformations being affected by salt [**Supplementary Fig. 9** and **11**].
